# Inevitability of the emergence and persistence of genetic parasites caused by thermodynamic instability of parasite-free states

**DOI:** 10.1101/182410

**Authors:** Eugene V. Koonin, Yuri I. Wolf, Mikhail I. Katsnelson

## Abstract

Genetic parasites, including viruses and mobile genetic elements, are ubiquitous among cellular life forms, and moreover, are the most abundant biological entities on earth that harbor the bulk of the genetic diversity. Here we examine simple thought experiments to demonstrate that both the emergence of parasites in simple replicator systems and their persistence in evolving life forms are inevitable because the putative parasite-free states are thermodynamically unstable.

## Background

Nearly all cellular life forms are hosts to various types of genetic parasites that exploit functional systems of the host cells to replicate their own genomes [1]. Only some bacteria with highly reduced genomes that themselves lead a symbiotic or parasitic lifestyle seem to lack genetic parasites that undoubtedly have been lost during the reductive evolution of these bacteria from free-living ancestors. Genetic parasites include viruses, transposons, plasmids and other semi-autonomous genetic elements (SAGE) [2] that display a broad range of relationships with the hosts, from acute antagonism, whereby a virus rapidly kills the host, to symbiosis when SAGE are not costly to the host and could even have beneficial effects [3, 4].

Strikingly, virus particles appear to be the most common biological entities on earth. In most environments, the ratio between virus particles and cells varies between 10 and 100 [4-7]. This enormous physical abundance of viruses is matched by vast genetic diversity so that most of the gene repertoire of the biosphere appears to be concentrated in viruses, even as exact number remain a matter of debate [8-10]. The prevalence of viruses in the biosphere is also paralleled by the abundance of SAGE integrated in genomes of cellular life forms. Integrated SAGE are present in virtually all genomes of cellular organisms (again, missing only in some intracellular symbionts and parasites), and in genomes of multicellular eukaryotes, SAGE-derived sequences quantitatively dominate the genome, comprising at least 50% of the DNA in vertebrates and up to 90% in plants [11]. Recruitment of sequences from SAGE for cellular functions is a common phenomenon that made substantial contributions to the evolution of cellular life forms [12-14].

The entire course of the evolution of life is a history of host-parasite co-evolution [15-17]. Being subject to the constant onslaught of genetic parasites, cellular life forms have evolved a plethora of defense mechanisms. A typical organism harbors and interacts with multiple types of genetic parasites (e.g. viruses, different families of transposons, and plasmids) which it holds at bay thanks to multiple defense strategies that include parasite exclusion, innate immunity and adaptive immunity [18-23]. The SAGE respond with counter-defense mechanisms that range from simple mutational escape from defense to dedicated multigene systems that specifically inactivate host defense systems. Notably, defense systems and SAGE including their counter-defense machineries are tightly linked in evolution. Enzymes involved in the mobility of SAGE, in particular, transposons are often recruited by host defense systems for roles in parasite genome inactivation and other functions, and conversely, SAGE recruit components of defense systems that then evolve to become agents of counter-defense [24-26].

Thus, the arms race, along with cooperation, between genetic parasites and their hosts are perennial features of the evolution of life. Why is this the case? Why do the parasites emerge in the first place? And, could some cellular organisms actually get rid of the parasites through highly efficient defense systems? Empirically, the answer to the latter question seems to be negative. Conceivably, the general cause of the inability of the hosts to eliminate the genetic parasites is the unescapable cost of maintaining sufficiently powerful defense systems [27-30]. Analysis of theoretical models of parasite propagation suggests that an important source of this cost, perhaps the primary one in microbes, could be that efficient anti-parasite defense has the side effect of curtailing horizontal gene transfer (HGT), which is an essential process in microbial evolution that allows microbes to avoid deterioration via Muller’s ratchet [31, 32]. Another major factor could be the effectively unavoidable autoimmunity [29, 33, 34]. However, what about the first, arguably, the most fundamental question: why do genetic parasites evolve to begin with? Again, empirically, there is a strong impression that the emergence of such parasites is inevitable. Not only are they ubiquitous in cellular life forms but they also evolve in various computer simulations of replicator system evolution [35-39]. Furthermore, it appears intuitive: genetic parasites can be considered cheaters, in game-theoretical terms, and as soon as, in a replicator system, there is a distributable resource, such as a replicase, cheaters would emerge to steal that resource without producing their share of it [40]. These, however, are informal considerations. Here we ask the question: is it possible to develop a theoretical framework that would allow a formal demonstration of the inevitability of the emergence of genetic parasites in evolving replicator systems, or else, that parasite-free replicator systems are after all possible?

### Thermodynamic instability of parasite-protected replicators

Let us try, as a gedunken experiment, to construct a self-replicating entity that is strictly resistant to parasites. Consider a simple system consisting of a replicator, serving as a template for itself, and the replicase it encodes (Figure 1). The replicator is assumed to contain the replicase-encoding signal (RES) (the replicase could be a protein, a ribozyme, under the RNA World model, or, in theory, any other entity capable of catalyzing replication of a template) and the replicase recognition signal (RRS).

**Figure 1.**
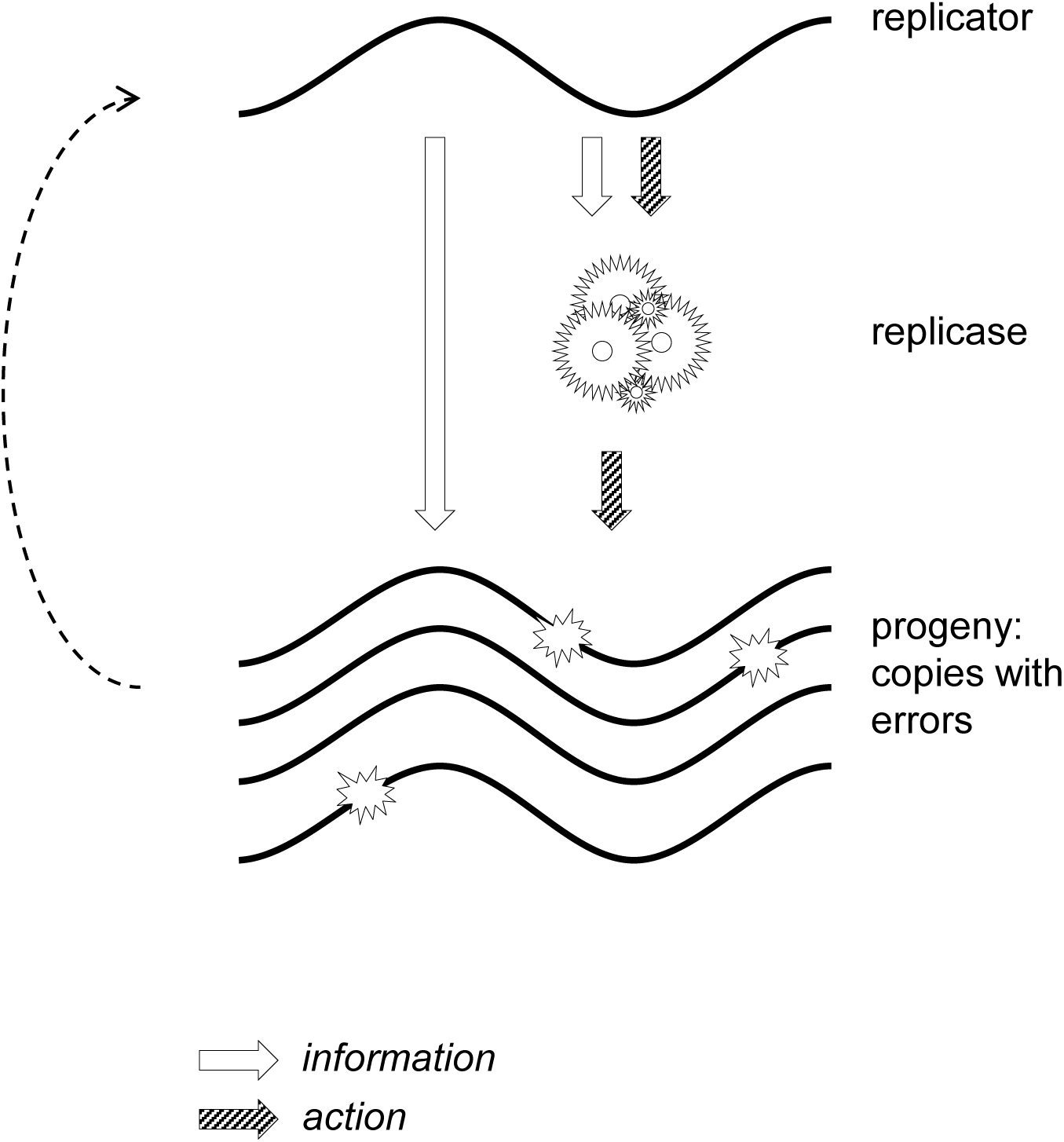
A replicator-replicase systems with heredity, variability and differential reproduction. The dotted arrow denotes differential reproduction of the copies of the original replicator that carry mutations.

Evolvability is a fundamental and inescapable property of such a replicator-replicase system [41]. To be evolvable, a system must possess three basic properties: 1) heredity (whereby the location of progeny in the phenotype space is correlated with that of the parents), 2) variability (whereby the progeny is not identical to the parents), and 3) differential reproduction (whereby the capability of a replicator to leave progeny is part of the phenotype). Heredity is ensured by replication with fidelity above the error catastrophe threshold. The replicator theory that was developed primarily by Eigen and colleagues demonstrates that, under simple fitness landscapes, there exists a replication fidelity threshold, below which the master sequence in a population of replicators cannot be efficiently passed across generations, so that the entire population collapses [42, 43]. Elucidation of the molecular mechanisms of primordial replication that could provide for crossing the error catastrophe threshold remains a daunting task that is central to the entire origin of life field. However, for the purpose of the present discussion, we assume that a sustainable replicator system with a minimally acceptable fidelity has evolved.

Variability is ensured because, at any temperature above 0 K, any process is subject to entropy-increasing fluctuations and, therefore, replication is inherently error-prone, under the second and third Laws of Thermodynamics.

Differential reproduction ensues from the fact that the replicator encodes the replicase that, in turn, copies the replicator itself. Mutations in both RES and RRS can affect the efficiency of replication.

If the resources that are available to the system are limited (i.e. the system does not support unlimited growth of all possible constituent parts), competition between individual replicators ensues and selection arises. In a system with finite memory storage, all information exchange, transfer and utilization processes carry memory clearing cost of at least *kT*ln2 J/bit, where *k* is Boltzmann constant and *T* is temperature (the existence and value of this minimum information cost is known as Landauer’s principle that is a corollary of the Second Law of Thermodynamics); in all known systems, this cost is many orders of magnitude higher [44-46]. Therefore, selection for cost reduction acts not only on the constituent parts of the system, but also on the information transfer processes themselves, effectively ensuring an upper limit on the fidelity of information recognition.

Selection acts on both RES (eliminating replicators encoding inefficient replicases) and RRS (eliminating replicators that are inefficient as templates, e.g. are poor replicase-binders), but these two selection processes act on the replicator through physically different agents (the replicator-encoded replicase and the replicator itself, respectively).

The dual nature of the replicator (acting as both the template and, directly or indirectly, as the replicase) necessitates that the information embedded in the RES and RRS is realized via physically different processes. The RES guides the formation of the replicase which, in turn, recognizes the RRS. Such recognition implies comparing the RRS in the replicator with some form of memory encoded in or attached to the replicase (Figure 2).

**Figure 2.**
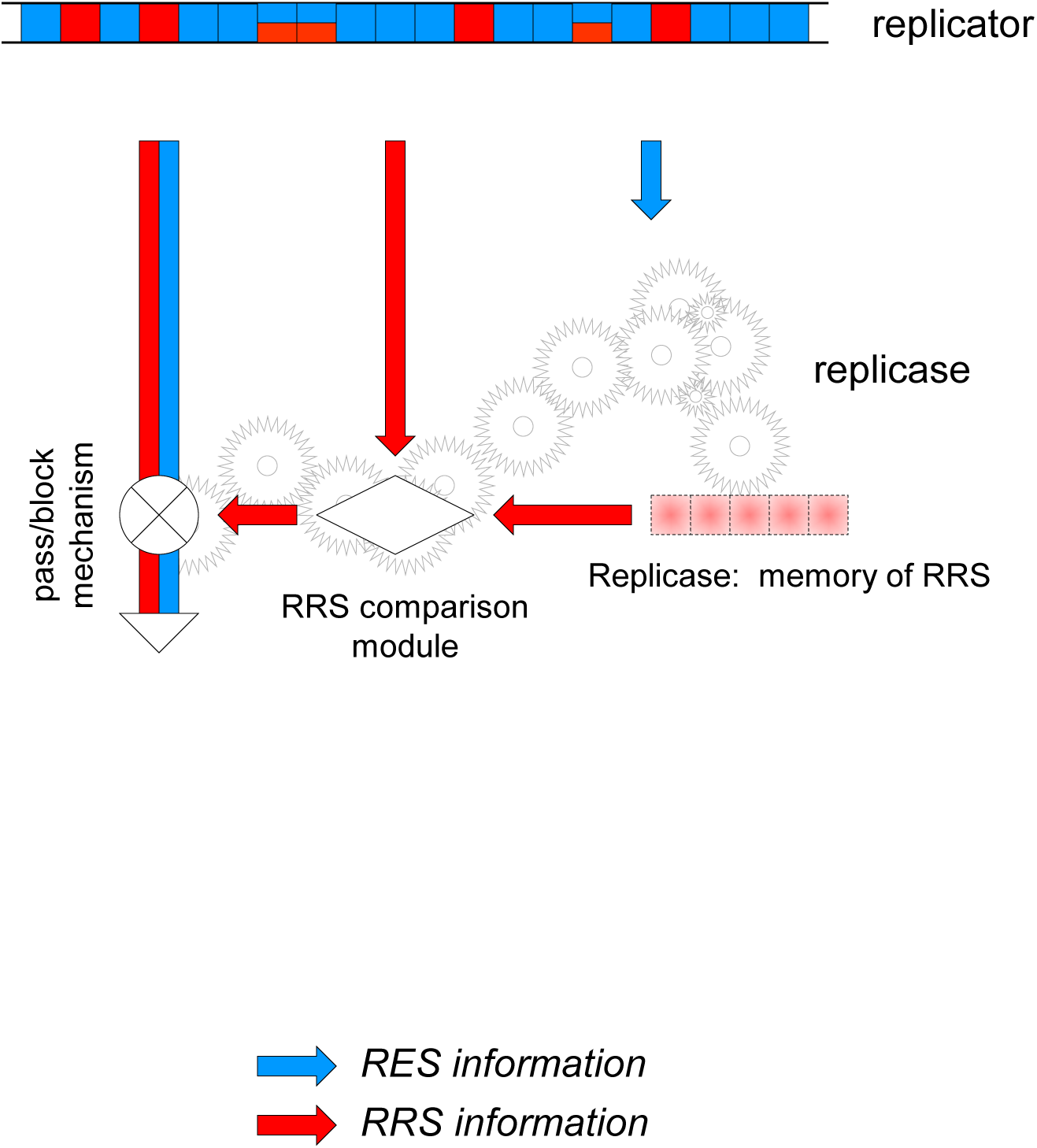
The replicase-encoding signal (RES) and replicase-recognition signal (RRS) in replicator-replicase systems. For generality, the RRS is shown as being distributed along the length of the replicator although in real genomes, this signal is often localized such that, for example, short terminal sequences are sufficient for the replication of a virus genome. The replicase structure carries memory of the RRS allowing recognition of competent templates (“pass/block mechanism”).

A general, simple way of parasite emergence involves skipping part of the RES during replication, resulting in a shorter replicator that consists of the RRS and, in the extreme, nothing else (Figure 3). This straightforward mechanism for parasite evolution is inspired by and is similar to the process of RNA shrinking that was observed during *in vitro* evolution in the classic early experiments of Spiegelman and colleagues [47-49].

**Figure 3.**
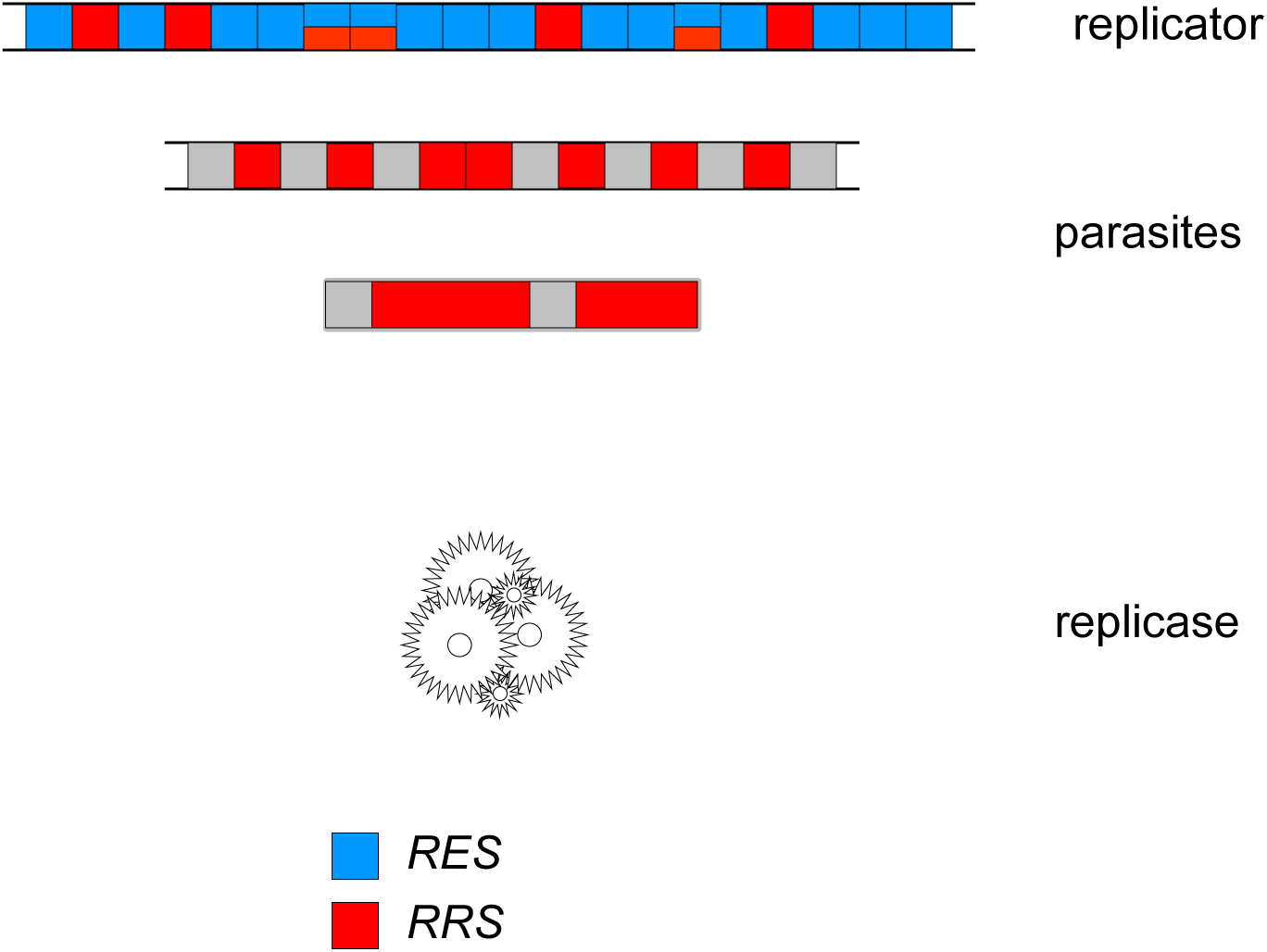
Emergence of parasites in replicator systems via deletion of portion of the RES.

Under the scheme in Figure 3, parasites emerge as long as the information content of the RRS is less than that in the full replicator, i.e. when the RRS is at least partially separable from the RES. If this is the case, a replicator containing the full RRS, but omitting at least some of the RES (RRS_p_ ≡ RRS, RES_p_ -> 0; the subscript ‘p’ denotes the respective signals in the parasite), would not only serve as a template as efficient as the original replicator, but would also enjoy an evolutionary advantage because replication of the smaller replicator is faster and requires less resources (building blocks, such as nucleotides, and energy). This makes the parasite-free equilibrium point of the replicator-parasite system unstable because deletion of any part of the RES yields more efficient replicators (Figure 4). Therefore, the system is vulnerable to parasite invasion, and moreover, such an invasion is inevitable under a non-zero parasite emergence rate (see Appendix for a more formal demonstration).

**Figure 4.**
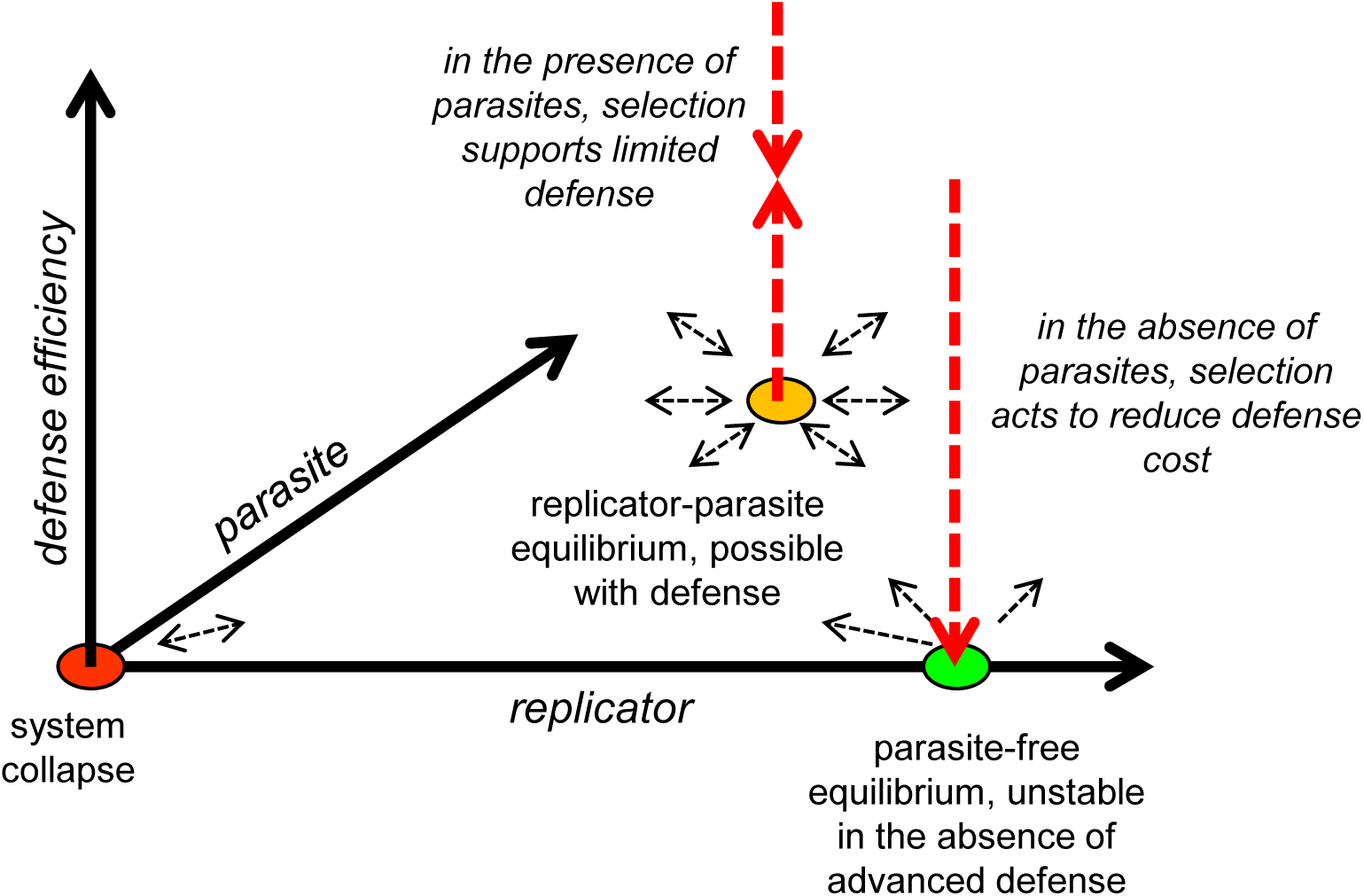
A conceptual phase diagram of the evolution of replicator systems.

It appears that, under this scheme, the only way to render the replicase-producing replicator parasite-protected is to make the RRS to include the entire RES (Figures 2 and 4). Such RRS ≡ RES configuration evidently rules out the emergence of a parasite because any mutation of the RES would also inactivate the RRS and prevent replication.

However, such a parasite-protected state is subject to the aforementioned instability (Figure 4). In the absence of parasites, perfect protection does not carry any benefits, but incurs a greater cost than less protected states. Given that the system is evolvable, an RRS < RES state will inevitably arise and outcompete the RRS ≡ RES progenitors that are, as shown above, prone to emergence of genetic parasites (RRS_p_ ≡ RRS, RES_p_ -> 0).

From a more abstract perspective, the fully protected RRS ≡ RES system corresponds to the maximally constrained, i.e. minimum entropy, state. The second law of thermodynamics effectively guarantees that it evolves into a higher entropy state, such that RRS < RES, and at least some parts of the RES can be mutated or deleted without compromising replication. The ensemble of higher entropy states is obviously more robust than the unique RRS ≡ RES state. In biological parlance, the higher entropy states are favored by selection via the ‘survival of the flattest’ route [50]. They will necessarily prevail because there are plenty of such states with similar fitness values, whereas the RRS ≡ RES state is singular. However, the problem with the ‘relaxed’ states of the replicator is that they are no longer protected from parasites because a parasite now can evolve that would exploit the RRS without producing the replicase (Figure 3). The Third Law of Thermodynamics dictates that the minimum (zero) entropy state can be stable only at 0 K. Under the detailed correspondence between thermodynamics and population genetics [51], the equivalent of temperature is the inverse effective population size, and accordingly, 0 K corresponds to an infinite population, which can exist only as an abstraction. Thus, in any realistic population, a parasite-free replicator system is inherently unstable and either goes extinct or rapidly spawns parasites (Figure 4). The latter case inevitably triggers the host-parasite arms race, whereby in the simplest case, the hosts evolve by selection for changed RRS allowing them to escape the parasites, whereas the parasites catch up. Furthermore, the competition also occurs between the parasites themselves and could eventually result in the emergence of ultimate parasites, those that consist entirely of the RRS. Such (near) ultimate parasites were the end result of Spiegelman’s experiments [49] and also exist in nature, namely the viroids, small, non-coding parasitic RNAs that cause disease in plants and rely entirely on a host-derived replication machinery [52, 53].

It should be noted that, because, at least in the simplest replicator system, the replication rate is inversely proportional to the genome length, the parasite have an intrinsic advantage in the arms race. In the well-mixed case, the preferential replication of parasites drives the host to extinction which, obviously, results in the collapse of the entire system (no replicase is produced anymore). However, compartmentalization can stabilize host-parasite systems. Thus, parasites, drive evolution of biological complexity [37-39, 54].

There seems to be a symmetry between the hypothetical, minimum entropy, parasite-resistant replicator and the ultimate parasite with RRS_p_ ≡ RRS, RES_p_ ≡ 0, another minimum entropy state, this one being the theoretical end result of the competition between parasites (Figure 3). As a minimum entropy state, the ultimate parasite cannot be evolutionarily stable either. The gedunken experiment described above certainly is an idealization. Realistically, when the minimum entropy state relaxes, the host-parasite arms race takes more complex forms. Most parasites are far from this ultimate state but rather possess a number of genes and encode a variety of functions. This complexity of parasites has to do with two strategies that parasites evolve to maximize their evolutionary success, namely: 1) overcoming the defense systems which the hosts evolve under the pressure for resistance to the parasites, and 2) surviving outside the host and disseminating among hosts [4]. The question, then, emerges: even though the above analysis shows that evolution of parasites in simple replicator systems is inevitable, is there a chance that evolution of defense systems would exterminate parasites?

### Costs and compromises of anti-parasite defense

The thought experiment described above also answers the question whether a perfect defense system can exist. A perfect self vs non-self discrimination, whereby a replicator possesses the means to reject or destroy any potential cheater, that is, any sequence other than a perfect copy of itself, is nothing but the same parasite-protected system with recognition based on the complete information on the self, i.e. RRS ≡ RES. We have already shown above that such a system is evolutionary unstable from pure thermodynamic considerations because it provides no benefit in the absence of parasites, and will inevitably devolve to an RRS < RES configuration (“leaky” defense).

The existence of an unavoidable cost implies that maintenance of any form of defense is subject to a cost-benefit tradeoff. Notably, a recent quantitative assessment of the selection coefficients (a measure of fitness cost) associated with different classes of genes in microbial genomes has shown that defense systems are as costly as the more benign SAGEs, such as transposons [30]. This result seems to be a genome-scale reflection of the intrinsic costliness of defense dictated by thermodynamics. In the actual biological context, this cost can be manifested in different forms, such as autoimmunity or interference of defense systems with HGT. The curtailment of HGT, in particular, could be a key factor that makes defense systems costly and causes their repeated loss [32]. However, at the bottom of it all seems to be the thermodynamic cost of information. Certainly, defense is not the only process associated with an information cost. Such cost is intrinsic to all processes of information transmission including replication, transcription, translation, as well as signal transduction. However, loss of genes encoding those other functions is often strongly deleterious to the organism, resulting in positive mean selection coefficients for the respective classes of genes [30]. This is not the case for defense systems because, of which the mean selection coefficient values for the defense genes are negative [30], and in accord with this observation, defense genes are lost in the course of evolution significantly more often than genes of other functional categories [55]. Due to these fitness costs, evolving organisms cannot build up defense systems to the extent that is required to eliminate parasites (Figure 4).

### A quasi-formal demonstration of the inevitability of the emergence and persistence of parasites

In the above, we demonstrate that a parasite-protected state, one in which RRS≡RES, is thermodynamically unstable. The inevitability of the emergence of parasites and their subsequent persistence follows from this demonstration. To present the argument succinctly and quasi-formally:

(1) The two signals that are essential in a replicator system, RRS and RES, are encoded in different genomic sequences.

(2) Thus, at least some parts of the RES can be deleted without inactivating the RRS. Hence genetic parasites emerge.

(3) The shorter the genome sequence of a genetic element the more efficient its replication is. Hence parasites accumulate in a replicator system and may bring it to collapse in a well-mixed case.

(4) A perfect defense system should, in the least, be able to recognize parasitic elements, i.e. detect missing parts of the host genome. At the end of the replication cycle, this information should be deleted. Under the Landauer principle, the cost of memory cleaning is at least *kTn*ln2 J (*n* bits distinguish the host and the parasite). This cost makes defense systems evolutionary unstable and precludes elimination of parasites.

## Conclusions

The problem of the ubiquity and persistence of genomic parasites throughout the evolution of life can be broken into two parts: i) emergence of parasites in primitive replicator systems, ii) persistence of parasites in evolving organisms. Our analysis of the basic aspects of host-parasite coevolution presented here suggests that there are fundamental thermodynamic causes of the inevitability of parasites on both stages. To put it most succinctly, both the hypothetical parasite-protected state in a simple replicator system and the putative secondary parasite-free state resulting from efficient action of evolved defense systems are thermodynamically unstable and can only exist transiently. These conclusions certainly are compatible with a wealth of observational data indicating that genetic parasites, along with defense systems, are enormously abundant in the biosphere and accompany virtually every cellular life form. The results of numerous mathematical and computational modeling studies on replicator evolution, in which parasites invariably appear, lead to the same conclusion. The simple thought experiments described here that start, effectively, from first principles emphasize the growing understanding that emergence as well as persistence of genetic parasites is an inalienable feature of evolving replicators and, as such, one of the central principles of biology.

## Appendix A.

Consider a simple system in which both the replicator and a parasite comprise single entities and the replicator entity can also function as the replicase (e.g., both the replicator and the parasite are single RNA molecules, and the replicator molecule in the native folded state can function as the replicase ribozyme). Replication of the replicator entity is governed by the second-order kinetics, where a template replicator meets an (identical) replicase to make a copy of the template. There is also a natural decay of both types of entities, occurring with the first-order kinetics. In an environment with a fixed carrying capacity (determined, e.g., by the influx of consumable resources), such a system behaves as a classical logistic model, with one exception: there exists a critical number of entities, below which the population is unsustainable due to the (second-order) replication lagging behind the (first-order) decay.

Likewise, a parasite entity replicates upon meeting a replicator (acting as a replicase) and decays spontaneously. The parasite population is subject to similar environmental restrictions as that of the replicator due to the limitation by the same resources.

The more general model that we consider here also includes the existence of a (potentially costly) defense system:

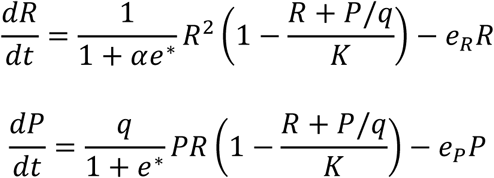

where *R* and *P* are the concentrations of the replicator and the parasite particles, respectively, *eR* 277 and *eP* are the corresponding decay rates and *K* is the environment carrying capacity (the replicator growth rate is taken to be 1 without loss of generality). The parasite enjoys an evolutionary advantage that is manifested in two ways: it both replicates faster and consumes fewer resources by the same factor *q* (the simple conceptual model of this effect is based on an RNA molecule that is shorter than the replicator by a factor of *q*). The parasite replication is countered by the defense systems that are embedded in the replicase and discriminate against the parasites with an efficiency *e*^*^. The action of the defense system is costly to the replicator, with the cost coefficient α (according to the Landauer principle, α > 0)

Upon introduction of the parasite to the population of the replicator near the equilibrium (*P* → 0, *dR*/*dt* → 0), the host defense prevents the parasite invasion (*dP*/*dt* < 0) only if

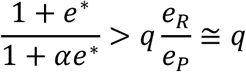

or, in other words, only if the effect of defense on the invading parasite relative to the cost of defense to the host is greater than the parasite advantage *q* (assuming comparable decay rates).

Obviously, the replicator population lacking the defense system (*e*^*^ = 0) cannot resist the parasite invasion. A defense system combining high efficiency with low cost is capable of protecting the host population, but the maintenance of such defense depends on regular invasion of parasites (otherwise, the evolutionary disadvantage due to the cost of defense would drive the defense system to extinction). In other words, a parasite-protected state of replicator system is unstable.

## Declarations

### Acknowledgements

Not applicable.

### Funding

EVK and YIW are supported by intramural funds of the US Department of Health and Human Services (to the National Library of Medicine).

### Availability of supporting data

Not applicable.

### Author contributions

EVK conceived of the project; EVK, YIW and MIK developed the theory and wrote the manuscript that was read and approved by all authors.

### Author’s information

Eugene V. Koonin and Yuir I. Wolf are at the National Center for Biotechnology Information, National Library of Medicine, National Institutes of Health, Bethesda, MD 20894, USA; Mikhail I. Katsnelson is at the Institute for Molecules and Materials, Radboud University, Nijmegen, 6525AJ, Netherlands.

### Competing interests

The authors declare that they have no competing interests.

### Consent for publication

Not applicable.

### Ethics approval and consent to participate

Not applicable.

